# ABCC1/MRP1 exports cGAMP and modulates cGAS-dependent immunity

**DOI:** 10.1101/2021.12.03.470980

**Authors:** Joanna H. Maltbaek, Jessica M. Snyder, Daniel B. Stetson

**Affiliations:** Departments of Immunology and Medicine, University of Washington School of Medicine; Seattle, WA 98109, USA; Department of Comparative Medicine, University of Washington School of Medicine, Seattle, WA 98195, USA

## Abstract

The DNA sensor cyclic GMP-AMP synthase (cGAS) is important for antiviral and anti-tumor immunity. cGAS generates cyclic GMP-AMP (cGAMP), a diffusible cyclic dinucleotide that activates the antiviral response through the adapter protein Stimulator of Interferon Genes (STING). cGAMP is negatively charged and cannot passively cross cell membranes, but recent advances have established a role for extracellular cGAMP as an “immunotransmitter” that can be imported into cells. However, the mechanism by which cGAMP exits cells remains unknown. Here, we identify ABCC1/MRP1 as an ATP-dependent cGAMP exporter that influences STING signaling and type I interferon production. We demonstrate that ABCC1 deficiency exacerbates cGAS-dependent autoimmunity in the *Trex1*^*-/-*^ mouse model of Aicardi-Goutières syndrome. These studies identify ABCC1-mediated cGAMP export as a key regulatory mechanism of the cGAS-STING pathway.

---

The cGAS-cGAMP-STING DNA sensing pathway has emerged as a key innate immune response that is important for antiviral immunity (*1*), contributes to autoimmune diseases (*2*), and mediates aspects of antitumor immunity (*3*). Cyclic GMP-AMP synthase (cGAS) is a nucleotidyltransferase enzyme that binds to double-stranded DNA and catalyzes the formation of cyclic GMP-AMP (cGAMP) (*4, 5*), a diffusible cyclic dinucleotide (CDN) that activates the adaptor protein stimulator of interferon genes (STING) on the endoplasmic reticulum (*6*). Activated STING then serves as a platform for the inducible recruitment of TANK-binding kinase 1 (TBK1), which phosphorylates and activates the transcription factor interferon regulatory factor 3 (IRF3), leading to the induction of the type I interferon (IFN)-mediated antiviral response (*7*).

The formation of cGAMP by cGAS has its roots in ancient nucleotidyltransferases that regulate signaling and metabolism in prokaryotes (*8, 9*). Like all RNA nucleotide-based second messengers that exist across all kingdoms of life (*10*), cGAMP is unable to passively enter or exit cells because of its two negatively charged phosphate groups that prevent movement across hydrophobic phospholipid bilayers that make up cellular membranes. Interestingly, the only known enzyme that degrades cGAMP in metazoans is an extracellular phosphodiesterase called ectonucleotide pyrophosphatase/phosphodiesterase 1 (ENPP1) (*11*), which hydrolyzes cGAMP into GMP and AMP (*12*). cGAMP can avoid ENPP1-mediated degradation via movement through gap junctions (*13*) and packaging into enveloped viral particles (*14, 15*). However, tumor cells have been shown to release cGAMP (*16*), and this pool of extracellular cGAMP is subject to potent degradation by ENPP1 (*17*). Moreover, extracellular cGAMP can be taken up by various importer channels and activate STING signaling in distant cells (*18-23*).

However, a fundamental question remains: how does cGAMP exit the cells that produce it? Here, we describe the ATP-binding cassette transporter ABCC1/MRP1 as a channel that mediates cGAMP export. We demonstrate that ABCC1-mediated cGAMP export is important for regulation of antiviral responses and control of cGAS-dependent autoimmunity.

## Results

### cGAMP is exported from live cells

Intracellular DNA detection activates both the cGAS-cGAMP-STING pathway to trigger IFN-mediated antiviral responses and the AIM2 inflammasome to activate rapid, inflammatory cell death known as pyroptosis (*24, 25*). This AIM2-dependent cell death pathway is particularly potent in myeloid cells, although it can function in non-myeloid cell types (*26*). To study cGAMP dynamics in the absence of the AIM2 inflammasome, we prepared primary bone marrow derived macrophages (BMMs) from mice lacking all 13 mouse AIM2-like receptors (ALRs) (*27*). As previously reported, we found that DNA-activated rapid cell death was almost completely abolished in *ALR*^*-/-*^ BMMs (Fig. 1A). We then measured intracellular and extracellular concentrations of cGAMP 8 hours after transfection with calf thymus (CT) DNA using a sensitive cGAMP ELISA (*28, 29*). We normalized the input volumes of cell extracts and extracellular media to allow direct comparison of absolute cGAMP amounts between these two compartments. We found that *ALR*^*-/-*^ BMMs produced significantly more total cGAMP than WT control BMMs, consistent with the enhanced IFN responses in AIM2- and ALR-deficient mice (Fig. 1B) (*24, 27*). However, we recovered the majority of this cGAMP from the extracellular media, not from inside the cells (Fig. 1B). Finally, *ALR*^*-/-*^*Sting*^*-/-*^ double knockout (DKO) BMMs exported similar amounts of cGAMP compared to *ALR*^*-/-*^ BMMs (Fig. 1B), demonstrating that STING signaling was not required for cGAMP export.

**Fig. 1.**
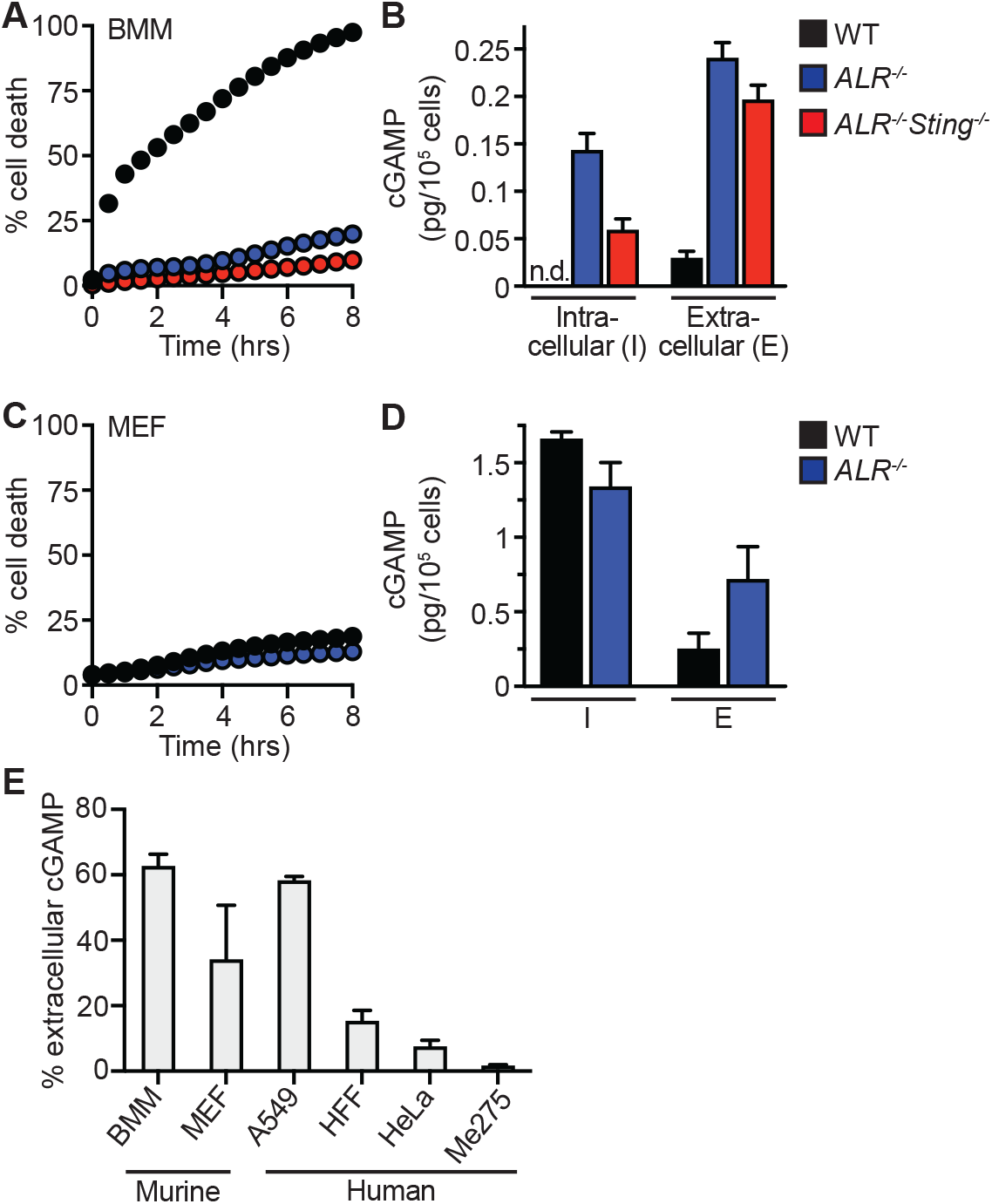
cGAMP is exported from live cells following cGAS activation. (**A**) Cell death of BMMs from WT, *ALR*^-/-^, *ALR*^-/-^*Sting*^-/-^ mice transfected with calf thymus DNA (CT DNA) quantified using an IncuCyte imaging system. (**B**) cGAMP quantification using ELISA from cell lysates and supernatants in (A) 8 hours after transfection. (**C**) Cell death of MEFs from WT or *ALR*^-/-^ mice transfected with calf thymus DNA (CT-DNA) quantified using an IncuCyte imaging system. (**D**) cGAMP quantification using ELISA from cell lysates and supernatants in (C) 8 hours after transfection. (**E**) Percent extracellular cGAMP of indicated cell types 8 hours after CT DNA transfection. cGAMP ELISA was used to determine relative extracellular and intracellular concentrations before calculating the percent extracellular. Error bars represent mean ± SD. All data are representative of three independent experiments.

We next tested primary murine embryonic fibroblasts (MEFs), which do not undergo ALR-dependent pyroptosis in response to DNA detection (Fig. 1C) (*27*). We found that MEFs exported significant amounts of cGAMP in the absence of cell death (Fig. 1D), but not as much as BMMs in relative extracellular versus intracellular cGAMP amounts (Fig. 1B). We extended these findings to immortalized human foreskin fibroblasts (HFF) and human tumor cell lines, including A549 lung adenocarcinoma cells (*30*), HeLa cervical carcinoma (*31*), and Me275 melanoma cells (*32*). We first verified that none of these cells expressed detectable levels of *Enpp1* mRNA or ENPP1 protein (fig. S1A, S1B). We found that the amount of extracellular cGAMP was low or nearly absent in HFF, HeLa, and Me275 cells, even though these cells all produced detectable intracellular cGAMP in response to DNA transfection (fig. S1C). A549 cells were the only human cell type tested that exported cGAMP at levels comparable to mouse BMMs (Fig. 1E). Thus, we have found that live cells export cGAMP, and we have identified a nearly 60-fold range of export “efficiency” among diverse mouse and human cell types (Fig. 1E).

### Identification of a drug that blocks cGAMP export

We considered the mechanisms by which cGAMP could be exported from live cells. Previous studies have shown that cGAMP can be enclosed in extracellular vesicles (*14, 15*), and a more recent study demonstrated that cGAMP can be exported as a soluble factor (*17*). We therefore focused on transmembrane channels as potential cGAMP exporters. There are two broad classes of such proteins: 1) channels that passively export substrates based on electrochemical gradients, and 2) channels that use the energy of ATP hydrolysis to move substrates across membranes, sometimes against a concentration gradient. We focused on active transport mechanisms that are accomplished by the ATP-binding cassette (ABC) family of transmembrane channels (*33*), guided by the prior identification of bacterial energy-dependent export channels that mediate efflux of cyclic dinucleotides (*8*). Human ABC transporters are named because of a shared intracellular domain that binds ATP and translates the energy of ATP hydrolysis into conformational changes that move cargo from the cytosol to extracellular space or into membrane-bound intracellular compartments. The human genome encodes 49 ABC transporters that move a broad spectrum of small molecules with diverse masses and chemical properties (*34*). Many ABC transporters have been scrutinized because of their ability to export chemotherapeutic drugs (*35, 36*), which has resulted in the development of a rich resource of pharmacological inhibitors of entire families of ABC channels. We used three such inhibitors of distinct classes of ABC transporters and measured their effect on cGAMP export from *ALR*^*-/-*^ BMMs: verapamil, which inhibits the ABCB1 transporter (*37*); KO-143, which blocks the ABCG2 transporter (*38*); and MK-571, a drug developed as an ABCC1 inhibitor (*39*). We found that verapamil did not influence cGAMP export, but pretreatment with MK-571 resulted in a significant, dose-dependent increase in intracellular cGAMP after DNA transfection (Fig. 2A), consistent with blockade of cGAMP export. The highest dose of KO-143 also significantly increased intracellular cGAMP, but it was previously reported that KO-143 has an inhibitory effect on the transport activity of both ABCB1 and ABCC1 at this concentration (*40*).

**Fig. 2.**
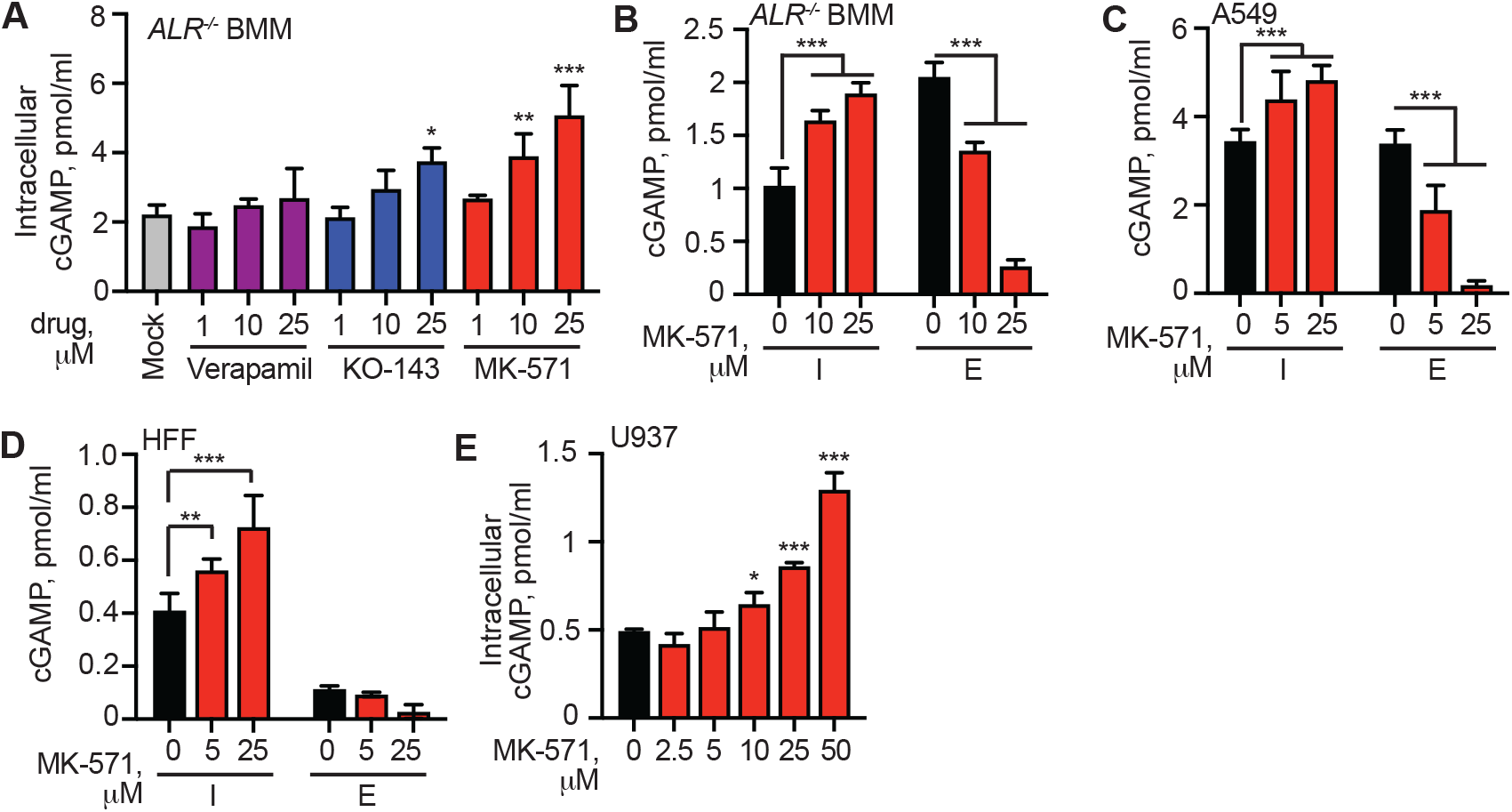
MK-571 blocks cGAMP export in mouse and human cells. (**A**) *ALR*^-/-^ BMMs were treated with 1, 10, or 25 µM of indicated inhibitor or mock, followed by CT DNA transfection. 8 hours later, intracellular cGAMP was quantified in cell lysates using ELISA. (**B**) *ALR*^-/-^ BMMs were treated with 10 or 25 µM MK-571 or mock, followed by CT DNA transfection. 8 hours later, cell lysates and supernatants were harvested and cGAMP was quantified using ELISA (I: intracellular; E: extracellular). (**C-E**) Various human cell types were treated with MK-571 (0-50 µM) followed by CT DNA transfection. 8 hours later, cell lysates and supernatants were harvested and cGAMP was quantified using ELISA. Statistical analysis was performed using a one-way ANOVA comparing all drug treatments to mock (A, E) or a two-way ANOVA comparing drug treatments to mock-treated conditions in the intracellular or extracellular compartment (B- D). Tests corrected for multiple comparisons using the Holm-Sidak method. Error bars represent mean ± SD. **p<0.01, ***p<0.001. Data are representative of two (A, E) or three (B-D) independent experiments.

Thus, we focused on the dose-dependent blockade of cGAMP export by MK-571. MK-571 pretreatment not only resulted in enhanced retention but also reduced export of cGAMP from BMMs (Fig. 2B). We extended these findings to human cells including A549 (Fig. 2C), HFF (Fig. 2D), and the human monocyte cell line U937 (*41*) (Fig. 2E). As with mouse cells, pre-treatment with MK-571 resulted in enhanced retention of cGAMP after DNA transfection in each human cell type. Furthermore, A549 cells treated with MK-571 had reduced cGAMP export (Fig. 2C), though extracellular cGAMP levels were low in HFF (Fig. 2D) or undetectable in U937. These data demonstrate that MK-571 potently blocks cGAMP export in both mouse and human cells.

### ABCC1 is a cGAMP exporter

MK-571 inhibits ABCC1, also known as multidrug resistance protein 1 (MRP1) (*42*), which has been demonstrated to mediate the export of cysteinyl LTC4, the conjugated estrogen E217βG, oxidized glutathione (GSH) (*43*), and sphingosine-1-phosphate (*44*). ABCC1 is a member of a family that includes several related transporters, and it is now known that MK-571 blocks other members of the ABCC/MRP family, including ABCC2/MRP2, ABCC4/MRP4, and ABCC5/MRP5 (*45, 46*). We developed quantitative RT-PCR assays to measure the expression of each member of the ABCC/MRP family. Moreover, to enable screening of potential candidate transporters using lentiCRISPR in primary BMMs, we crossed our *ALR*^-/-^ mice with mice that constitutively express Cas9 (*47*). We found that *ALR*^-/-^Cas9^+^ BMMs expressed detectable mRNA transcripts for five of the eight ABCC family members (Fig. 3A). We designed guide RNAs (gRNAs) to target these five expressed ABCC channels, cloned them into a lentiCRISPR vector, prepared lentivirus particles, and transduced primary BMMs followed by selection in puromycin (*27-29, 48*). As a control for transduction and for off-target effects, we included a gRNA targeting *Abcc6*, which was not expressed in BMMs (Fig. 3A). After selection, we observed significant but incomplete disruption of the target genes using TIDE analysis (Tracking of Indels by Decomposition) (*49*) (Fig. 3B). We found that disruption of *Abcc1*, but not any of the other expressed *Abcc* genes, resulted in a significant decrease in the percent of extracellular cGAMP recovered following DNA transfection (Fig. 3C, fig. S2A). We repeated this experiment, including an additional M1 non-targeting control gRNA, together with a second gRNA targeting a distinct site in the *Abcc1* gene. Both *Abcc1* gRNAs depleted ABCC1 protein levels (Fig. 3D) and decreased extracellular cGAMP (Fig. 3E, fig. S2B). To assess whether *Abcc1* is responsible for all cGAMP export in BMMs, we crossed ABCC1-deficient mice to our *ALR*^-/-^ mice. Heterozygous *Abcc1*^*+/-*^*ALR*^-/-^ BMMs had reduced ABCC1 protein levels, whereas homozygous *Abcc1*^-/-^*ALR*^*-/-*^ BMMs lacked ABCC1 protein expression (Fig. 3F). We found a significant reduction in extracellular cGAMP after DNA stimulation in both *Abcc1*^*+/-*^ and *Abcc1*^*-/-*^ BMMs compared to controls (Fig. 3G, fig. S2C). Pretreatment with MK-571 further reduced cGAMP export (Fig. 3G, fig. S2C). These data demonstrate that ABCC1 is responsible for most cGAMP export in BMMs, and that these cells have additional transport mechanism(s) that are sensitive to MK-571.

**Fig. 3.**
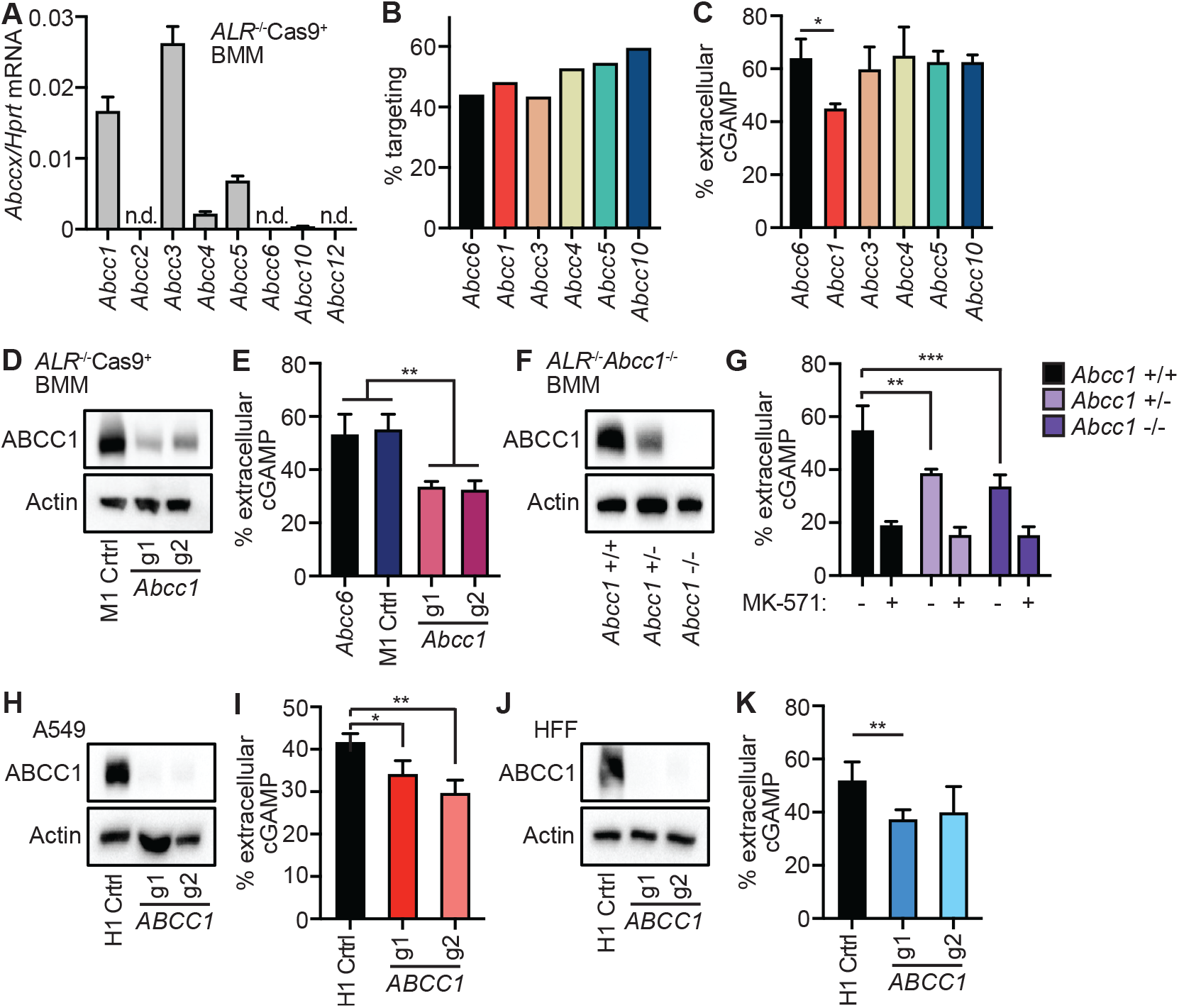
ABCC1 is a cGAMP exporter. (**A**) Quantification of ABCC family member mRNA transcript expression in *ALR*^-/-^Cas9^+^ BMMs by RT-qPCR. (**B**) *ALR*^-/-^Cas9^+^ BMMs were transduced with lentiCRISPR encoding the indicated ABCC family-specific gRNAs, selected for 3 days, and then percent genomic targeting was calculated using Sanger sequencing and Tracking of Indels by DEcomposition (TIDE) analysis. (**C**) Cells from (B) were transfected with CT DNA and then 8 hours later cGAMP was quantified in supernatants and cell lysates using ELISA. (**D**) *ALR*^-/-^Cas9^+^ BMMs were transduced with lentiCRISPR encoding *Abcc1* or M1 control-specific gRNAs as described in (B) followed by Western blot analysis for ABCC1 protein. (**E**) Cells from (D) were transfected with CT DNA and then 8 hours later cGAMP was quantified in supernatants and cell lysates using ELISA. (**F**) BMMs were harvested from *ALR*^-/-^ BMMs mice that were crossed to different ABCC1 genotypes and then evaluated by Western blot for ABCC1 protein. (**G**) Cells from (F) were transfected with CT DNA and then 8 hours later cGAMP was quantified in supernatants and cell lysates using ELISA. (**H**) A549 cells were transduced with lentiCRISPR encoding *ABCC1*- or H1 control-specific gRNAs as described in (B) and ABCC1 protein expression was evaluated by Western blot. (**I**) Cells from (H) were transfected with CT DNA and then 8 hours later cGAMP was quantified in supernatants and cell lysates using ELISA. (**J**) HFFs were transduced with lentiCRISPR encoding *ABCC1*- or H1 control-specific gRNAs as described in (B) and ABCC1 protein was assessed by Western blot. (**K**) Cells from (J) were transfected with CT DNA and then 8 hours later cGAMP was quantified in supernatants and cell lysates using ELISA. Statistical analysis was performed using a one-way ANOVA comparing targeted lines to relevant controls and corrected for multiple comparisons using the Holm-Sidak method. Error bars represent mean ± SD. *p<0.05, **p<0.01, ***p<0.001. Data are representative of two (A-C, F, G) or three (D, E, H-K) independent experiments.

Next, we tested whether disruption of *ABCC1* reduces cGAMP export in human cells. We designed two distinct gRNAs to target human *ABCC1* with lentiCRISPR as described above. We first tested these gRNAs in A549s and found that they depleted ABCC1 protein (Fig. 3H). We assessed functional loss of ABCC1 transport activity following targeting by staining cells with Fluo-3, a calcium-binding fluorescent compound that is a known substrate of ABCC1-dependent export (*50, 51*). We used measurement of Fluo-3 fluorescence as a proxy for ABCC1 activity: cells with high ABCC1 activity are dimmer than cells with low ABCC1 activity (fig. S2D). Flow cytometry analysis showed that *ABCC1* targeted A549 cells had a significant increase in Fluo-3 mean fluorescence intensity (MFI) compared to H1 control targeted cells, demonstrating that these gRNAs reduce ABCC1 transport activity (fig. S2E, S2F). *ABCC1* targeted A549 cells had a significant decrease in percentage of extracellular cGAMP compared to control H1 targeted cells (Fig. 3I, fig. S2G). We also used these gRNAs to target *ABCC1* in HFFs. While both *ABCC1* gRNAs depleted ABCC1 protein in HFFs (Fig. 3J) and reduced the percent of extracellular cGAMP (Fig. 3K, fig. S2H), only one gRNA resulted in a significant decrease in the percent of extracellular cGAMP compared to H1 control targeted cells. Taken together, these data demonstrate that disruption of human *ABCC1* reduces cGAMP export. Moreover, and similar to *Abcc1*^*-/-*^ mouse cells (Fig. 2), additional cGAMP export mechanism(s) exist in human cells that cannot be explained by ABCC1.

### ABCC1 overexpression enhances cGAMP export

We found that ABCC1 protein expression levels correlated well with the diverse efficiencies of cGAMP export that we identified among several mouse and human cell lines (Fig. 4A, 4B). We therefore tested whether ABCC1 overexpression could increase cGAMP export in cells with lower endogenous ABCC1 expression. To do this, we transduced HFFs, which had relatively low ABCC1 expression and poor cGAMP export efficiency, with lentivirus encoding human ABCC1. To test the requirement for ATP hydrolysis in cGAMP export, we introduced a K1333M point mutation into the second nucleotide binding domain (NBD) of ABCC1 that is known to reduce ABCC1-mediated LTC4 transport by approximately 70% (*52*). After selection of transduced cells, we observed increased ABCC1 protein levels (Fig. 4C) and decreased Fluo-3 retention in the ABCC1-overexpressing HFFs relative to those transduced with control lentivirus (Fig. 4D, 4E), demonstrating increased ABCC1 function. Surprisingly, HFFs expressing K1333M mutant ABCC1 also had a significantly lower Fluo-3 MFI than control HFFs, but not as low as HFFs overexpressing WT ABCC1 (Fig. 4D, 4E). To our knowledge, Fluo-3 fluorescence has never been studied in the context of the K1333M mutation, so it is possible that Fluo-3 export by ABCC1 is partially ATP-independent. We next tested these cells for cGAMP export after DNA transfection. We found that HFFs overexpressing WT ABCC1 exported more than triple the percent of extracellular cGAMP compared to controls (Fig. 4F, fig. S3A). In contrast, the K1333M mutant failed to enhance cGAMP export (Fig. 4F, fig. S3A).

**Fig. 4.**
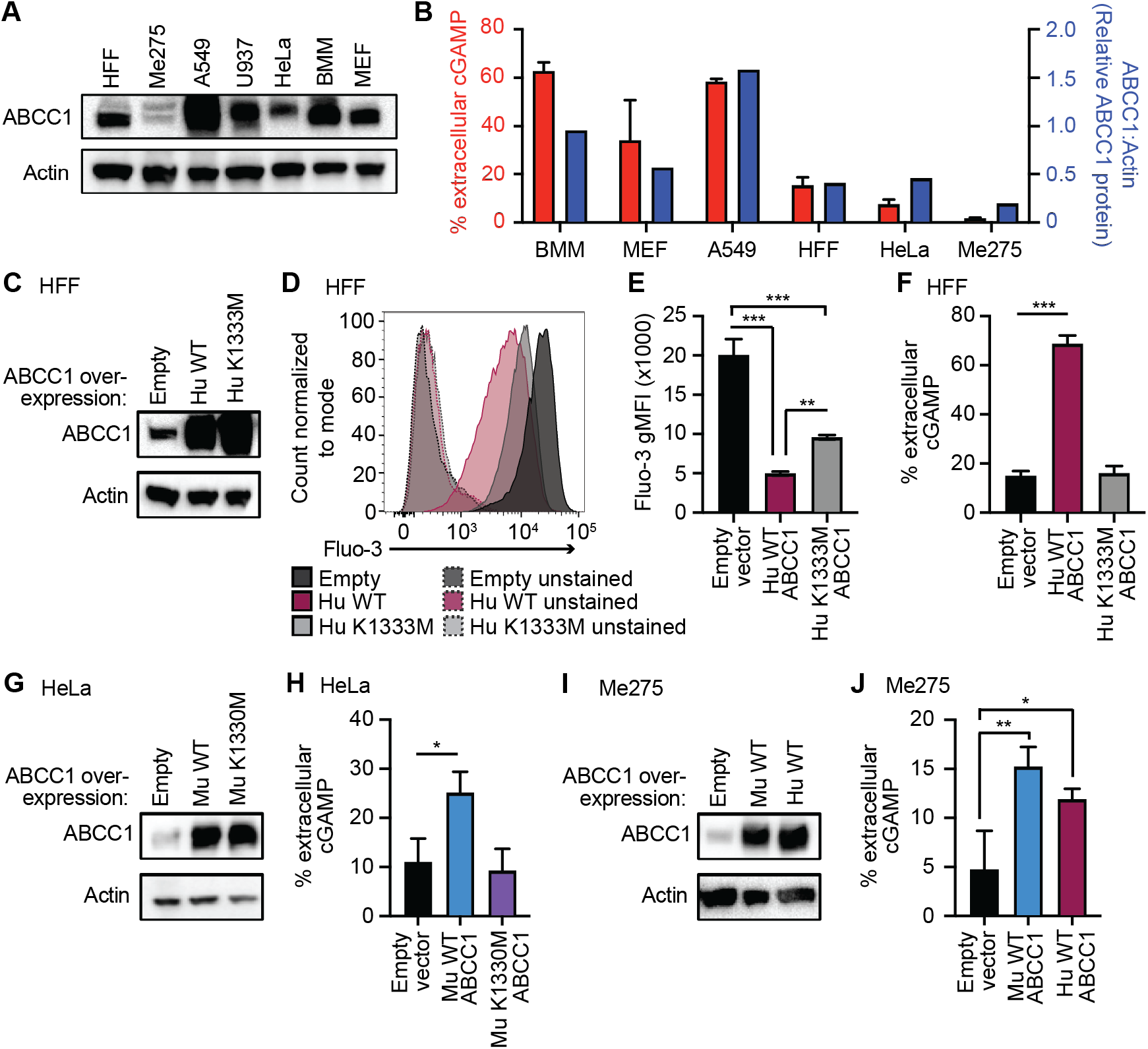
ABCC1 overexpression enhances cGAMP export. (**A**) Western blot analysis of indicated cells for ABCC1 protein expression. (**B**) cGAMP export efficiency calculated in Fig. 1E overlayed with densitometry from (A) for ABCC1 protein expression normalized to Actin expression. (**C**) HFFs were transduced with lentivirus encoding human WT or K1333M mutant ABCC1 or control (empty vector) and selected for 5 days in hygromycin. Cells were evaluated for ABCC1 protein expression by Western blot. (**D**) Cells from (C) were stained with Fluo-3AM and incubated for 1 hour, followed by flow cytometric quantification of Fluo-3AM staining intensity. (**E**) Quantification of gMFI from (D). (**F**) Cells from (C) were transfected with CT DNA and then 8 hours later cGAMP was quantified in supernatants and cell lysates using ELISA. (**G**) HeLa cells were transduced with lentivirus encoding murine WT or K1330M mutant ABCC1 or control (empty vector) and selected for 5 days in hygromycin. Cells were evaluated for ABCC1 protein expression by Western blot. (**H**) Cells from (G) were transfected with CT DNA and then 8 hours later cGAMP was quantified in supernatants and cell lysates using ELISA. (**I**) Me275 cells were transduced with lentivirus encoding human or murine WT ABCC1 or control (empty vector) and selected for 5 days in hygromycin. Cells were evaluated for ABCC1 protein expression by Western blot. (**J**) Cells from (I) were transfected with CT DNA and then 8 hours later cGAMP was quantified in supernatants and cell lysates using ELISA. Statistical analysis was performed using a one-way ANOVA comparing WT or mutant ABCC1-overexpression cells to empty vector control (D, F, H) or comparing the means of all groups to each other (C) and corrected for multiple comparisons using the Holm-Sidak method. Error bars represent mean ± SD. *p<0.05, **p<0.01, ***p<0.001. All data are representative of three independent experiments.

To evaluate functional orthology between human and mouse ABCC1, we cloned the murine ABCC1 open reading frame into a lentiviral vector, as well as the corresponding K1330M NBD mutation (*52*). We transduced HeLa cells, which, like HFFs, have relatively low endogenous ABCC1 expression and low efficiency of cGAMP export. After selection of transduced cells, we observed increased ABCC1 protein in HeLa cells expressing both the WT ABCC1 and the K1330M mutant relative to those transduced with control lentivirus (Fig. 4G). We transfected these cells with CT DNA and found that HeLa cells overexpressing WT murine ABCC1, but not K1330M ABCC1, had a significant increase in the percent of extracellular cGAMP (Fig. 4H, fig. S3B). Finally, we found that overexpression of either human or mouse ABCC1 significantly enhanced cGAMP export in Me275 human melanoma cells (Fig. 4I, 4J, fig. S3C), which were the poorest cGAMP exporters among the cell types we examined. Together, these data demonstrate that both human and mouse ABCC1 mediate ATP-dependent cGAMP export.

### Modulation of cGAMP export influences cell-intrinsic STING signaling

cGAMP binding to STING activates TBK1- and IRF3-dependent transcription of type I IFN genes (*7*). We hypothesized that reduction of intracellular cGAMP concentrations through ABCC1-dependent export would similarly reduce cell-intrinsic STING signaling. Consistent with this hypothesis, we found that pretreatment of HFFs with MK-571 potently enhanced DNA-activated *IFNB1* transcription but not RNA-activated *IFNB1* transcription (Fig. 5A). This was accompanied by increased phosphorylated IRF3 protein but not phosphorylated STING protein (fig. S4). Similarly, MK-571 pretreatment enhanced the response in cells treated with extracellular cGAMP, as evidenced by enhanced phospho-STING and phospho-IRF3 protein (Fig. 5B), as well as increased *IFNB1* transcription (Fig. 5C). These data demonstrate that cGAMP export limits cell intrinsic STING signaling and enhances the stimulatory activity of imported cGAMP.

**Fig. 5.**
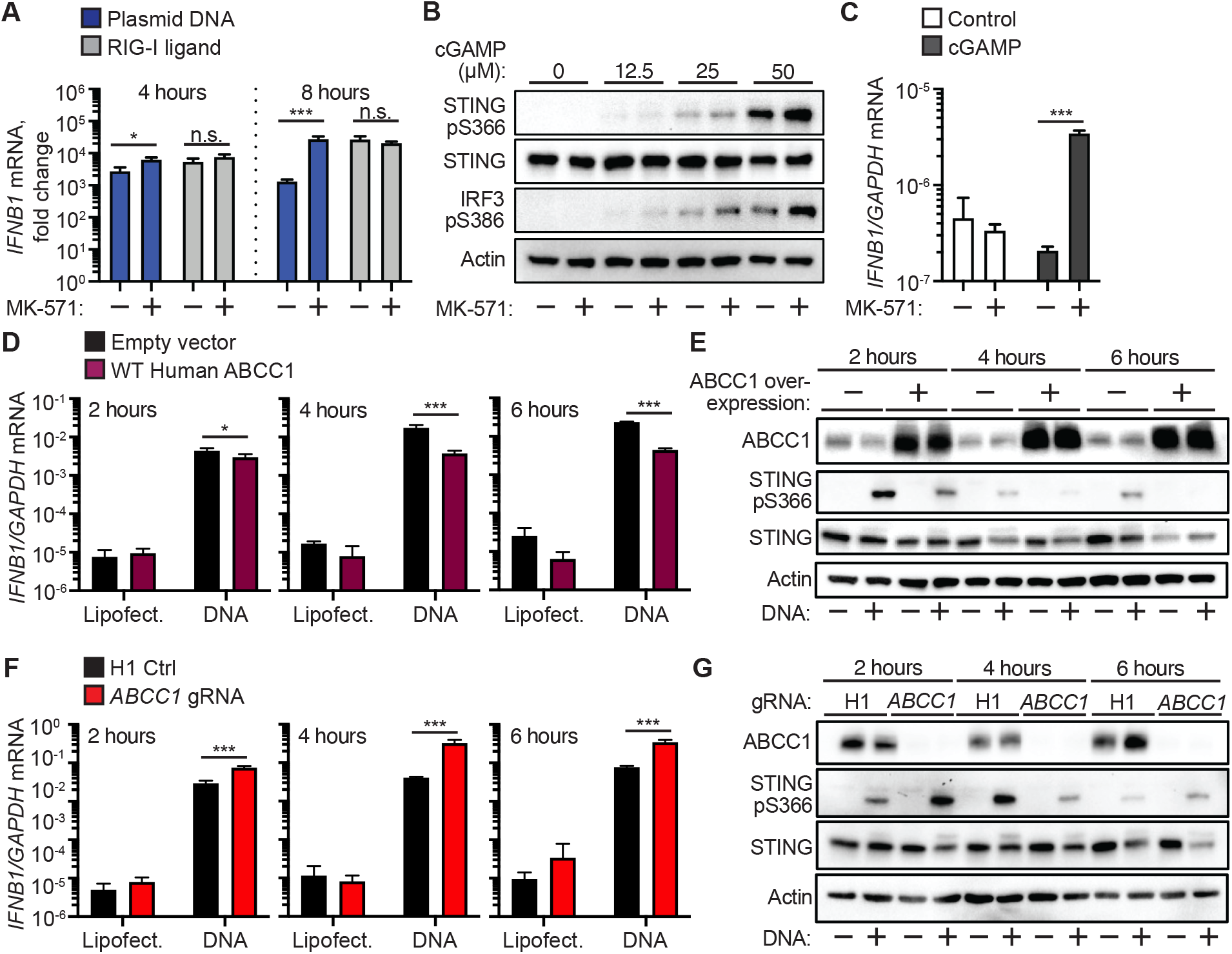
Modulation of cGAMP export influences cell-intrinsic STING signaling. (**A**) Quantification of *IFNB1* induction by RT-qPCR in HFFs that were treated with 25 µM MK-571 or mock followed by transfection with plasmid DNA or RIG-I ligand for 4 or 8 hrs. (**B**) HFFs were treated with 25 µM MK-571 or mock, and then cGAMP (0-50 µM) was added to the extracellular media. Western blot analysis was performed 4 hours after cGAMP addition for phosphorylated STING, STING, and phosphorylated IRF3. (**C**) HFFs were treated with 25 µM MK-571 or mock, and then cGAMP (0 or 50 µM) was added to the extracellular media. *INFB1* induction was quantified by RT-qPCR 4 hours after cGAMP addition. (**D**) Quantification of *IFNB1* induction by RT-qPCR in HFFs overexpressing ABCC1 or empty vector control following transfection with CT DNA. Cells were harvested at the indicated time points. (**E**) Western blot analysis of cells from (D) for ABCC1, phosphorylated STING, and STING protein expression. (**F**) Quantification of *IFNB1* induction by RT-qPCR in *ABCC1*- or H1 control-targeted HFFs following transfection with CT DNA. Cells were harvested at the indicated time points. (**G**) Western blot analysis of cells from (F) for ABCC1, phosphorylated STING, and STING protein expression. Statistical analysis was performed using a two-way ANOVA comparing mock or MK-571 treatment within each transfected ligand group (A, C) or comparing control to ABCC1-modulated cells within each transfected ligand group (D, F). All tests were corrected for multiple comparisons using the Holm-Sidak method. Error bars represent mean ± SD. *p<0.05, ***p<0.001. All data are representative of three independent experiments.

Given our findings that pharmacological blockade of cGAMP export enhanced STING signaling, we hypothesized that modulation of ABCC1 expression alters Type I IFN production. To test this, we stimulated our ABCC1-overexpressing HFFs with CT DNA and found that these cells had significantly lower *IFNB1* transcript levels compared to control cells throughout a 6-hour time course (Fig. 5D). Within this same experiment, we found decreased phospho-STING at each time point in ABCC1-overexpressing cells, consistent with the reduction in Type I IFN levels (Fig. 5E). Conversely, we found that DNA-stimulated HFFs targeted for *ABCC1* with lentiCRISPR (Fig. 3J) had increased *IFNB1* transcript levels and accelerated kinetics of phospho-STING protein when compared to H1 non-targeting control cells (Fig. 5F, 5G). These findings demonstrate that modulation of ABCC1 expression alters the antiviral response to DNA and suggest that ABCC1 acts as a negative regulator of cell intrinsic STING signaling.

### ABCC1 deficiency enhances cGAS-dependent autoimmunity in Trex1^-/-^ mice

Mutations in the human gene that encodes the TREX1 DNA exonuclease cause a rare and severe autoimmune disease called Aicardi-Goutières Syndrome (AGS) (*53*). We have shown that TREX1 is an essential and specific negative regulator of the cGAS-cGAMP-STING DNA sensing pathway (*54*). *Trex1*^-/-^ mice have a median life span of around 110 days and develop severe autoimmunity that requires cGAS (*55, 56*), STING (*57*), type I IFNs (*54*), and lymphocytes (*54, 57*). Thus, we utilized *Trex1*^-/-^ mice to explore the contribution of cGAMP export in a clinically relevant mouse model of human autoimmunity. Since ABCC1 expression correlated negatively with type I IFN production after DNA stimulation (Fig. 5), we hypothesized that loss of ABCC1 in *Trex1*^-/-^ mice would lead to accelerated mortality from enhanced IFN-mediated disease. We intercrossed *Trex1*^-/-^ and *Abcc1*^-/-^ mice and monitored survival. We found that both *Abcc1*^-/-^*Trex1*^-/-^ and *Abcc1*^+/-^*Trex1*^-/-^ mice exhibited significantly accelerated mortality compared to *Abcc1*^+/+^*Trex1*^-/-^ mice (Fig. 6A), consistent with the similarly reduced cGAMP export in both *Abcc1*^*-/-*^ and *Abcc1*^*+/-*^ BMMs (Fig. 3G). At 35 days of age, *Abcc1*^-/-^*Trex1*^-/-^ mice were more severely runted than *Abcc1*^+/+^*Trex1*^-/-^ mice (Fig. 6B). We measured tissue cGAMP levels and found that hearts from over half of the *Abcc1*^*-/-*^ mice that we sampled had detectable levels of cGAMP, whereas we could not detect cGAMP in the hearts of any *Abcc1*^*+/+*^ (WT) or *Cgas*^*-/-*^ mice (Fig. 6C). This increase is consistent with previously reported accumulation of GSH (a distinct ABCC1 substrate) in hearts of *Abcc1*^*-/-*^ mice (*58*). Heart cGAMP levels were detectable in most *Abcc1*^*+/+*^*Trex1*^*-/-*^ mice and significantly increased in all *Abcc1*^*-/-*^*Trex1*^*-/-*^ mice relative to control mice. Despite the limit of detection of the cGAMP ELISA assay, these data suggest that ABCC1 deficiency results in increased tissue cGAMP levels in both steady state and during cGAS-dependent disease.

**Fig. 6.**
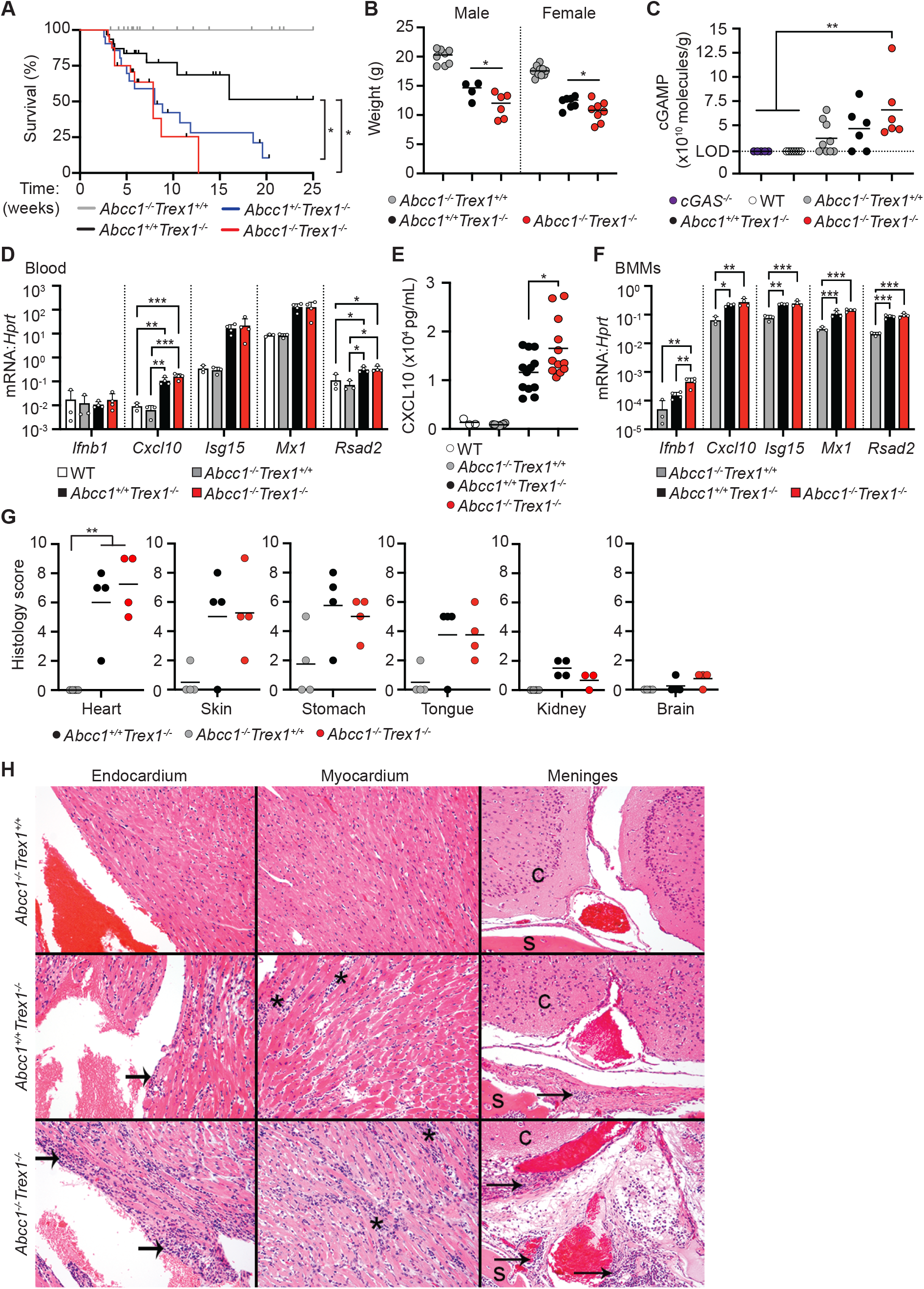
ABCC1 deficiency enhances cGAS-dependent autoimmunity in *Trex1*^*-/-*^ mice. (**A**) Survival of *Abcc1*^*-/-*^*Trex1*^*+/+*^ (n= 33), *Abcc1*^*+/+*^*Trex1*^*-/-*^ (n= 31), *Abcc1*^*+/-*^*Trex1*^*-/-*^ (n= 21), and *Abcc1*^*-/-*^*Trex1*^*-/-*^ (n= 24) mice. (**B**) Weights of mice at 35 days of age; *Abcc1*^*+/+*^*Trex1*^*-/-*^ (n= 8 M, 10 F), *Abcc1*^*+/-*^*Trex1*^*-/-*^ (n= 4 M, 7 F), and *Abcc1*^*-/-*^*Trex1*^*-/-*^ (n= 6 M, 8 F). (**C**) Quantification of intracellular cGAMP recovered from heart tissue by ELISA; *cGAS*^*-/-*^ (n= 5), WT (n= 7), *Abcc1*^*-/-*^ *Trex1*^*+/+*^ (n= 9), *Abcc1*^*+/+*^*Trex1*^*-/-*^ (n= 6), *Abcc1*^*-/-*^*Trex1*^*-/-*^ (n= 6). Values are normalized to individual heart weights (LOD = limit of detection at 2.3×10^10^ molecules/g). (**D**) Quantification of mRNA transcript levels by RT-qPCR in blood of the indicated genotypes; n= 3-4 mice per group. (**E**) Quantification of Cxcl10 protein in serum measured by ELISA of the indicated genotypes; WT (n= 3), *Abcc1*^*-/-*^ (n= 8), *Abcc1*^*+/+*^*Trex1*^*-/-*^ (n= 12), *Abcc1*^*-/-*^*Trex1*^*-/-*^ (n= 12). (**F**) Quantification of mRNA transcript levels by RT-qPCR in BMMs of the indicated genotypes; n= 3 mice per group. (**G**) Histological scores in the indicated tissues measured at 40 days of age; n= 4 mice per group. (**H**) Representative (of 4 of each genotype) H&E-stained heart and brain tissues sections from *Abcc1*^*-/-*^*Trex1*^*+/+*^, *Abcc1*^*+/+*^*Trex1*^*-/-*^, and *Abcc1*^*-/-*^*Trex1*^*-/-*^ mice from (G). Statistical analysis for the survival curves was calculated with a log-rank (Mantel-Cox) test (A). Statistical analysis for all other experiments was performed using a one-way ANOVA comparing each group to every other group (B, C, F), comparing transcript level of each gene between genotypes (D, E), or comparing *Abcc1*^*-/-*^*Trex1*^*+/+*^ to *Abcc1*^*+/+*^*Trex1*^*-/-*^ and *Abcc1*^*-/-*^*Trex1*^*-/-*^ mice for each tissue examined (G). All ANOVA tests were corrected for multiple comparisons using the Holm-Sidak method. Error bars represent mean ± SD. *p<0.05, **p<0.01, ***p<0.001.

We evaluated the expression levels of interferon stimulated genes (ISGs), and we found more significantly enhanced expression of *Cxcl10* mRNA in peripheral blood from *Abcc1*^-/-^*Trex1*^-/-^ mice than *Abcc1*^+/+^*Trex1*^-/-^ mice when compared to WT and *Abcc1*^*-/-*^ controls (Fig. 6D). Other ISGs were similarly elevated in both *Abcc1*^-/-^*Trex1*^-/-^ mice and *Abcc1*^+/+^*Trex1*^-/-^ mice (Fig. 6D). Consistent with this, we found that serum CXCL10 protein levels were significantly elevated in *Abcc1*^-/-^*Trex1*^-/-^ mice compared to *Abcc1*^+/+^*Trex1*^-/-^ mice (Fig. 6E). We next prepared BMMs from age-and sex-matched mice and found that expression of *Ifnb1* mRNA was significantly increased in *Abcc1*^-/-^*Trex1*^-/-^ BMMs compared to *Abcc1*^+/+^*Trex1*^-/-^ BMMs, whereas the expression of several ISGs was similarly increased in both *Trex1*^-/-^ genotypes relative to *Abcc1*^*-/-*^ controls (Fig. 6F).

Lastly, we performed a blinded histological analysis of affected tissues, comparing *Abcc1*^-/-^*Trex1*^-/-^ mice and *Abcc1*^+/+^*Trex1*^-/-^ mice to *Abcc1*^-/-^ controls. We detected similarly severe inflammation in *Abcc1*^-/-^*Trex1*^-/-^ mice and *Abcc1*^+/+^*Trex1*^-/-^ mice, with the heart tissue most severely affected in both genotypes (Fig. 6G, 6H). We also noted the presence of focal to multifocal, generally mild, perivascular lymphoid aggregates (with lower numbers of other inflammatory cells) associated with the meninges and periosteum of the skull in 3/4 *Abcc1*^-/-^ *Trex1*^-/-^ mice compared to 1/4 *Abcc1*^+/+^*Trex1*^-/-^ mice (Figure 6H). Taken together, these data demonstrate that ABCC1 is an essential negative regulator of the cGAS-cGAMP-STING pathway *in vivo* in a model of chronic cGAS activation.

## Discussion

In this study, we identify a mammalian cGAMP exporter, the ATP-binding cassette protein ABCC1. We show that loss of ABCC1 expression enhances intracellular cGAMP concentrations and, conversely, overexpression of ABCC1 reduces intracellular cGAMP concentrations. We further show that modulation of ABCC1 expression influences STING signaling and alters type I IFN production. Finally, we demonstrate that ABCC1 deficiency exacerbates cGAS-dependent autoimmunity *in vivo* in the well-characterized *Trex1*^*-/-*^ mouse model of human AGS.

Genetic disruption of *ABCC1* reduced but did not completely ablate cGAMP export *in vitro* (Fig. 3), suggesting that alternative cGAMP export mechanisms exist. This is perhaps unsurprising considering that multiple cGAMP importer channels have been identified to date (*18-23*). It was recently proposed that the LRRC8-containing volume regulated anion channel (VRAC) might serve as a “passive” cGAMP exporter, but these studies clearly demonstrated that LRRC8 can only export cGAMP under artificial hypotonic conditions that activate the channel through cell swelling (*20, 21*). While our CRISPR screen did not implicate other ABCC family members in cGAMP export (Fig. 3C), our finding that MK-571 further inhibits cGAMP export in *Abcc1*^*-/-*^ BMMs (Fig. 3G) and the fact that other nucleotide derivates such as cyclic AMP and cyclic GMP can be moved by ABCC family proteins (*59-61*) suggest that cGAMP might be a substrate of multiple ABC transporters. To identify additional cGAMP exporters, our ALR-deficient, Cas9-expressing mice remain a valuable tool for CRISPR-based screens in primary cells. Regardless of the potential for multiple cGAMP export mechanisms, we emphasize that ABCC1 plays a nonredundant role in cGAMP export and regulation of cGAS-STING signaling, both *in vitro* and *in vivo*.

Our findings have important implications for our understanding of the regulatory mechanisms involved in cGAS-STING signaling. We demonstrate that genetic ablation or pharmacological blockade of ABCC1 enhances cell-intrinsic STING signaling, whereas overexpression of ABCC1 reduces STING signaling. Thus, by limiting intracellular cGAMP concentrations, ABCC1 negatively regulates the cGAS-STING pathway and provides a mechanism to expose extracellular cGAMP to ENPP1-mediated degradation (*11*). In further support of this, loss of *Abcc1* in the *Trex1*^*-/-*^ mouse model led to accelerated and exacerbated disease (Fig. 6).

Our studies did not directly test the “immunotransmitter” function of cGAMP signaling that is additionally regulated by cGAMP degradation and cGAMP import. Ultimately, the influence of cGAMP export on STING signaling could be context and disease dependent, and it is possible that ABCC1-mediated export can play a positive regulatory role in propagating STING signaling to bystander cells. For example, we identified highly variable cGAMP export efficiencies across multiple human cancer cell lines (Fig. 2) and found that ABCC1 overexpression converted poor cGAMP exporters into more efficient exporters (Fig. 4). Prior studies highlighted an essential role for tumor-derived extracellular cGAMP in priming antitumor immune responses (*16, 17*). Our findings raise the interesting possibility that high expression of ABCC1 on tumor cells enhances antitumor immune responses through increased extracellular cGAMP in the tumor microenvironment. However, it is known that overexpression of ABCC1 by tumor cells renders them resistant to certain chemotherapeutic drugs that have been previously defined as export substrates of ABCC1 (*43*). Thus, the potential clinical utility of ABCC1 inhibitors in cancer patients to limit efflux of anti-cancer chemotherapeutics might be offset by their potential to reduce the export of an important innate immune signal that primes anti-tumor immunity. Therefore, our finding that cGAMP is a substrate of ABCC1 warrants further study of the consequences of ABCC1 blockade in cancer.

Finally, there have been numerous recent advances in our understanding of the ancient evolutionary origins of the cGAS-STING pathway. These include the discoveries of diverse cGAS orthologs, the generation of 2’-5’-linked oligonucleotides, and the functional conservation of STING in prokaryotic antiviral immunity (*62-66*). Our identification of ABCC1 as a cGAMP exporter in mammals raises interesting comparisons to the regulation of intracellular CDN concentrations in prokaryotes. The ABC transporter superfamily exists not only in eukaryotes but also prokaryotes and archaea (*67*), and the regulation of intracellular CDN concentrations through energy-dependent efflux is documented in bacteria (*8*). We postulate that cGAMP export by ABCC1 fits into this framework in which the central regulatory components involved in mammalian cGAS-STING signaling all evolved from ancient mechanisms of defense against bacteriophages. A detailed evolutionary analysis is called for to explore this intriguing possibility. The possibility that ABCC1 exports other CDNs, such as those of microbial origin, also remains to be determined.

In summary, we have identified ABCC1 as an important molecular site of action for cGAMP export. Our discovery highlights ABCC1 as a negative regulator of cell-intrinsic STING signaling and completes the cycle of cGAMP production, export, and import, further rationalizing the existence of an extracellular cGAMP degradation mechanism. Further investigation into ABCC1-mediated cGAMP export has the potential to yield novel therapeutic approaches to enhance the protective functions of type I IFN in human diseases.

## Acknowledgments

We thank N. Mausolf and S. Miller for expert mouse colony management, B. Johnson (University of Washington Histology and Imaging Core) for histology preparation, and all the members of the Stetson lab for helpful discussions.

## Author contributions

Conceptualization: JHM, DBS

Formal analysis: JHM

Methodology: JHM, DBS

Investigation: JHM, JMS

Visualization: JHM, JMS

Funding acquisition: JHM, DBS

Writing – original draft: JHM

Writing – review & editing: JHM, JMS, DBS

## Data and materials availability

All data are available in the main text or the supplementary materials.

## Materials and Methods

### Study Design

The objective of this study was to define a mechanism of cGAMP export. To investigate this, we surveyed various inhibitors of ABC transporters for the ability to block cGAMP export. We screened the ABCC/MRP family of transporters for specific cGAMP exporters. Functional characterization of the candidate transporter was performed with CRISPR targeting, lentiviral overexpression systems, and a mouse model of human AGS. Technical and biological replicate information are included in the appropriate figure legends. Quantification of histology was performed blinded by a board-certified pathologist.

### Mice

*ALR*^*-/-*^, *Sting (Tmem173)*^*-/-*^, and *Trex1*^*-/-*^ mice have been described (*27, 55, 57*). C57BL/6J wild-type (stock #000664) and *Abcc1*^*-/-*^ (stock #028129) mice were purchased from Jackson Laboratories. All mice used in this study were C57BL/6J and housed in a specific pathogen-free facility at the University of Washington with approval of the University of Washington Institutional Animal Care and Use Committee.

### Cell lines and tissue culture

Primary mouse BMMs and MEFs were derived and cultured as previously described (*48*). HeLa, U937, A549, and HepG2 were purchased from ATCC. Me275 cells, established at the Ludwig Institute for Cancer Research (*32*), were provided by A. Rongvaux (Fred Hutchinson Cancer Research Center). Telomerase-immortalized human foreskin fibroblasts (HFFs) were provided by D. Galloway (Fred Hutchinson Cancer Research Center). All adherent cell lines were cultured in DMEM supplemented with 10% FCS, L-glutamine, penicillin/streptomycin, sodium pyruvate, and HEPES. U937 cells were grown in RPMI supplemented as above and differentiated prior to stimulation using 100 nM phorbol myristoyl acetate (PMA) for 24 hours and then rested in media for an additional 24 hours without PMA prior to treatment.

### Quantification of cell death

BMMs and MEFs were seeded at a density of 2×10^4^ and 1×10^4^, respectively, in a 24-well plate. Cell death was assayed with a 2-color IncuCyte Zoom in-incubator imaging system (Essen Biosciences, Ann Arbor, MI, USA) and analyzed as described (*68*). SytoxGreen and SytoGreen (25 nM, Life Technologies) were used to calculate the frequency of dead cells.

### cGAMP measurements

For *in vitro* measurements, extracellular cGAMP was measured directly from cell supernatants that were maintained at a volume of 200 µL. To quantify intracellular cGAMP, cells were washed once with PBS and then lysed in 200 µL RIPA buffer (150 mM NaCl, 1% Triton-X-100, 0.5% sodium deoxycholate, 0.1% SDS, 50 mM Tris pH 8.0) for 15 minutes on ice. Lysates were then cleared of insoluble material before being used for cGAMP measurement. Relative cGAMP measurements were quantified using Direct 2’3’-Cyclic GAMP ELISA Kit (Arbor Assays).

To quantify cGAMP in mouse heart tissues, hearts were cleaned and weighed, minced, and then digested in dissociation buffer (2.7 mg/mL Collagenase A (Sigma), 23 U/mL DNase I (Sigma), 2 mM CaCl2 in PBS) for 1 hour at 37° C. Digestion was terminated with termination buffer (2% FBS, 5 mM EDTA in PBS). Samples were strained through 70 µm mesh strainers, centrifuged, and then subjected to red blood cell lysis in buffer (150 mM NH4CL, 10 mM KHCO3, 0.1 mM Na EDTA in water). Remaining cells were washed twice and suspended in 200 µL RIPA buffer for 15 minutes of lysis, then centrifuged to clear of insoluble material before direct measurement in cGAMP ELISA Kit. Limit of detection (LOD) was calculated by averaging the weights of all control (*cGAS*^*-/-*^ and WT) hearts to use in calculation of molecules per gram of tissue with the given LOD of the ELISA Kit (0.04 pMol/mL).

### Cell treatments and stimulations

For all *in vitro* experiments, cells were seeded in triplicate at a density of 1×10^5^ (A549, HeLa, Me275), 2×10^5^ (U937), or 2.5×10^4^ (HFFs) cells per well in 24-well plates. For nucleic acid transfections, calf thymus genomic DNA (Sigma) was diluted in water and used at 4 µg/mL (for human cells) or 1 µg/mL (for mouse cells); RIG-I ligand was synthesized *in vitro* as previously described using HiScribe T7 High Yield RNA Synthesis Kit (*69*) and used at 1 µg/mL; midiprepped pcDNA3 was used for plasmid stimulations at 4 µg/mL. For all transfections, nucleic acids were complexed with Lipofectamine 2000 (Invitrogen) at a ratio of 1 µg nucleic acid to 1 µL lipid. 2’3’ cGAMP (Invivogen) was diluted in water and directly added to cell culture media at 12.5 – 50 µM. MK-571, Verapamil, and KO-143 (Sigma) were suspended in DMSO and used to treat cells at concentrations from 1-50 µM for 1 hour prior to stimulation. Mock-treated cells received the same amounts of plain DMSO.

### Western blotting and antibodies

Cells were harvested in RIPA lysis buffer supplemented with phosphatase and protease inhibitors (Pierce). Lysates were vortexed and incubated on ice for 15 minutes followed by clearing by centrifugation for 15 minutes. Proteins were separated on 4-12% Bis-Tris SDS-PAGE gel (Life Technologies) and transferred to Immobilon-P PVDF membrane (Millipore).

Blots were blocked in 5% BSA/TBST or 5% non-fat dry milk depending on primary antibody used. Membranes were probed overnight at 4° C with the following primary antibodies: anti-mouse/human ABCC1 (Abcam ab260038), anti-mouse/human Actin (Cell Signaling #3700), anti-human phospho-STING (Cell Signaling #19781), anti-mouse phospho-STING (Cell Signaling #72971), anti-mouse/human STING (Cell Signaling #13647), anti-human phospho-IRF3 (Abcam ab196035). Membranes were then probed with goat anti-rabbit-HRP or goat anti-mouse-HRP (Jackson ImmunoResearch) and developed by ECL 2 chemiluminescence (Pierce).

### Fluo-3 AM staining and flow cytometry

Single cell suspensions were washed in PBS followed by staining with 2.5 uM Fluo-3 AM (Sigma) in FACS buffer (5% FBS in HBSS) and allowed to incubate for 20 mins on ice. 4 volumes of FACS buffer were then added and samples were incubated for an additional 40 mins at 37 C without CO2. After washing, samples were acquired on the Canto-II (BD Bioscience). Raw data were analyzed with FlowJo v10.

### LentiCRISPR targeting

VSV-G pseudotyped, self-inactivating lentivirus was prepared by transfecting a 60-80% confluent 10-cm plate of HEK 293T cells with 1.5 μg of pVSV-G expression vector, 3 μg of pMDLg/pRRE, 3 μg pRSV-Rev and 6 μg of pRRL lentiCRISPR vectors using Poly(ethyleneimine) (PEI; Sigma). Media was replaced 24 hours post-transfection and harvested 24 hours later for filtration with a 0.45 μm filter (SteriFlip, Millipore). Approximately 1 million cells were transduced with 10 mL filtered virus. Following 3-5 days of selection in appropriate antibiotic, successful targeting was verified through population-level Sanger sequencing and analysis using Tracking of Indels by DEcomposition (TIDE) (*49*).

For CRISPR/Cas9 gene targeting, we generated pRRL lentiviral vectors in which a U6 promoter drives expression of a gRNA, and an MND promoter drives expression of Cas9, a T2A peptide, and either puromycin or hygromycin resistance (*27*). Primary mouse cells targeted in Fig. 3 received lentivirus that did not encode Cas9, as the cells came from mice that have constitutive Cas9 expression. gRNA sequences were designed using Benchling. gRNA sequences are as follows, where the (G) denotes a nucleotide added to enable robust transcription off the U6 promoter:

Murine M1 non-targeting control: GCGAGGTATTCGGCTCCGCG (*70*)

Murine *Abcc1* guide 1: (G)TCAGAACACGGTCCTCACAT

Murine *Abcc1* guide 2: (G)TGGGCTGACCAGTAACACTG

Murine *Abcc3:* (G)ACACCACAGCAGAACACCGA

Murine *Abcc4:* (G)TGCGAGCCAAGAAGGACTCG

Murine *Abcc5:* (G)TCCGGAACCTGGGTTCCTCG

Murine *Abcc6:* (G)TGAAGAGGTGGGACATCCGG

Murine *Abcc10:* (G)TACTAGGCACCGACTCCGAG

Human H1 non-targeting control: (G)ACGGAGGCTAAGCGTCGCAA (*70*)

Human *ABCC1* guide 1: (G)ATAGACAGCCCCAATGACAG

Human *ABCC1* guide 2: (G)ACGATGATGACGTCCACCTG

### Cloning of ABCC1

PCR and In-Fusion cloning (Takara Clontech) were used to generate ABCC1 constructs. WT and K1333M mutant human ABCC1 constructs were generated using a human ABCC1 cDNA clone (Transomic, Clone ID TOH6003) as template. Mouse WT and K13330 mutant ABCC1 constructs were generated using a mouse ABCC1 cDNA clone (Transomic, Clone ID BC090617) as template.

### Quantitative RT-PCR

For quantitative RT-PCR analysis, triplicate cell samples or 50 uL fresh mouse blood were harvested into Trizol reagent before purification via Direct-zol RNA miniprep (Zymo Research) per manufacturer’s instructions with an additional dry spin after disposing of the final wash to prevent carryover. cDNA was generated using EcoDry premix (Takara Bio). Samples were assayed in duplicate and transcript expression was measured using iTaq Universal SYBR green supermix (BioRad) on the Bio-Rad CFX96 Real-Time system.

Human gene PCR primer sequences are as follows:

*GAPDH* Fwd: 5’-AACAGCCTCAAGATCATCAGC-3’, *GAPDH* Rev: 5’- CACCACCTTCTTGATGTCATC-3’,

*IFNB1* Fwd: 5’-ACGCCGCATTGACCATCTATG-3’, *IFNB1* Rev: 5’- CGGAGGTAACCTGTAAGTCTGT-3’.

Mouse gene PCR primer sequences are as follows:

*Hprt* Fwd: 5’-GTTGGATACAGGCCAGATTTGTTG-3’, *Hprt* Rev: 5’- GAGGGTAGGCTGGCCTATAGGCT-3’,

*Abcc1* Fwd: 5’-CTCTATCTCTCCCGACATGACC-3’, *Abcc1* Rev: 5’- AGCAGACGATCCACAGCAAAA-3’,

*Abcc2* Fwd: 5’-CCCTGCTGTTCGATATACCAATC-3’, *Abcc2* Rev: 5’- TCGAGAGAATCCAGAATAGGGAC-3’,

*Abcc3* Fwd: 5’-CTGTGCACACAGAAAACCCG-3’, *Abcc3* Rev: 5’- GGACACCCAGGACCATCTTG-3’,

*Abcc4* Fwd: 5’-AGCTGAGAATGACGCACAGAA-3’, *Abcc4* Rev: 5’- ATATGGGCTGGATTACTTTGGC-3’,

*Abcc5* Fwd: 5’-AGTCCTGGGTATAGAAGTGTGAG-3’, *Abcc5* Rev: 5’- ATTCCAACGGTCGAGTTCTCC-3’,

*Abcc6* Fwd: 5’-AAGGAGGTACTAGGTGGGCTT-3’, *Abcc6* Rev: 5’- CCAGTAGGACCCTTCGAGC-3’,

*Abcc10* Fwd: 5’-CCTAGTGCTGACCGTGTTGT-3’, *Abcc10* Rev: 5’- TAGGTTGGCTGCAGTCTGTG-3’,

*Abcc12* Fwd: 5’-ATGATGCCGGGCTACTCTC-3’, *Abcc12* Rev: 5’- CAGGGTGTCTACGGTCAGC-3’,

*Ifnb* Fwd: 5’-GCACTGGGTGGAATGAGACTATTG-3’ *Ifnb* Rev: 5’- TTCTGAGGCATCAACTGACAGGTC -3’,

*Cxcl10* Fwd: 5’-AAGTGCTGCCGTCATTTTCTGCCTC-3’, *Cxcl10* Rev: 5’- CTTGATGGTCTTAGATTCCGGATTC-3’,

*Mx1* Fwd: 5’-GACCATAGGGGTCTTGACCAA-3’, *Mx1* Rev: 5’- AGACTTGCTCTTTCTGAAAAGCC-3’,

*Isg15* Fwd: 5’-GGTGTCCGTGACTAACTCCAT-3’, *Isg15* Rev: 5’- TGGAAAGGGTAAGACCGTCCT-3’.

### CXCL10 ELISA

Mouse blood was collected into Eppendorf tubes and allowed to clot for 45 minutes at room temperature, followed by two rounds of centrifugation and collection of top serum layer. Serum samples were assayed directly with Mouse IP-10 (CXCL10) ELISA Kit (Abcam) and analyzed per manufacturer’s instructions.

### Histology and Pathology

Tissues were fixed in 10% neutral buffered formalin and routinely paraffin embedded. Tissue sections (5 μm) were stained with hematoxylin and eosin and histological scores were assigned in a blinded manner as previously described (*57*) with the following modifications: histology scores for skin and subcutaneous muscles of the head were combined; scores for the kidney reflect both inflammation scores for the interstitium as well as the extent of pathology.

Images were captured from glass slides using NIS-Elements BR 3.2 64-bit and plated in Adobe Photoshop Elements. Image white balance, lighting, and contrast were adjusted using auto corrections applied to the entire image. Original magnification is stated.

### Experimental replicates and statistics

Quantitative data were visualized and analyzed using GraphPad Prism software, ImageJ, and FlowJo. Specific statistical tests and experimental replicate numbers are noted in the figure legends.

**Fig. S1.**
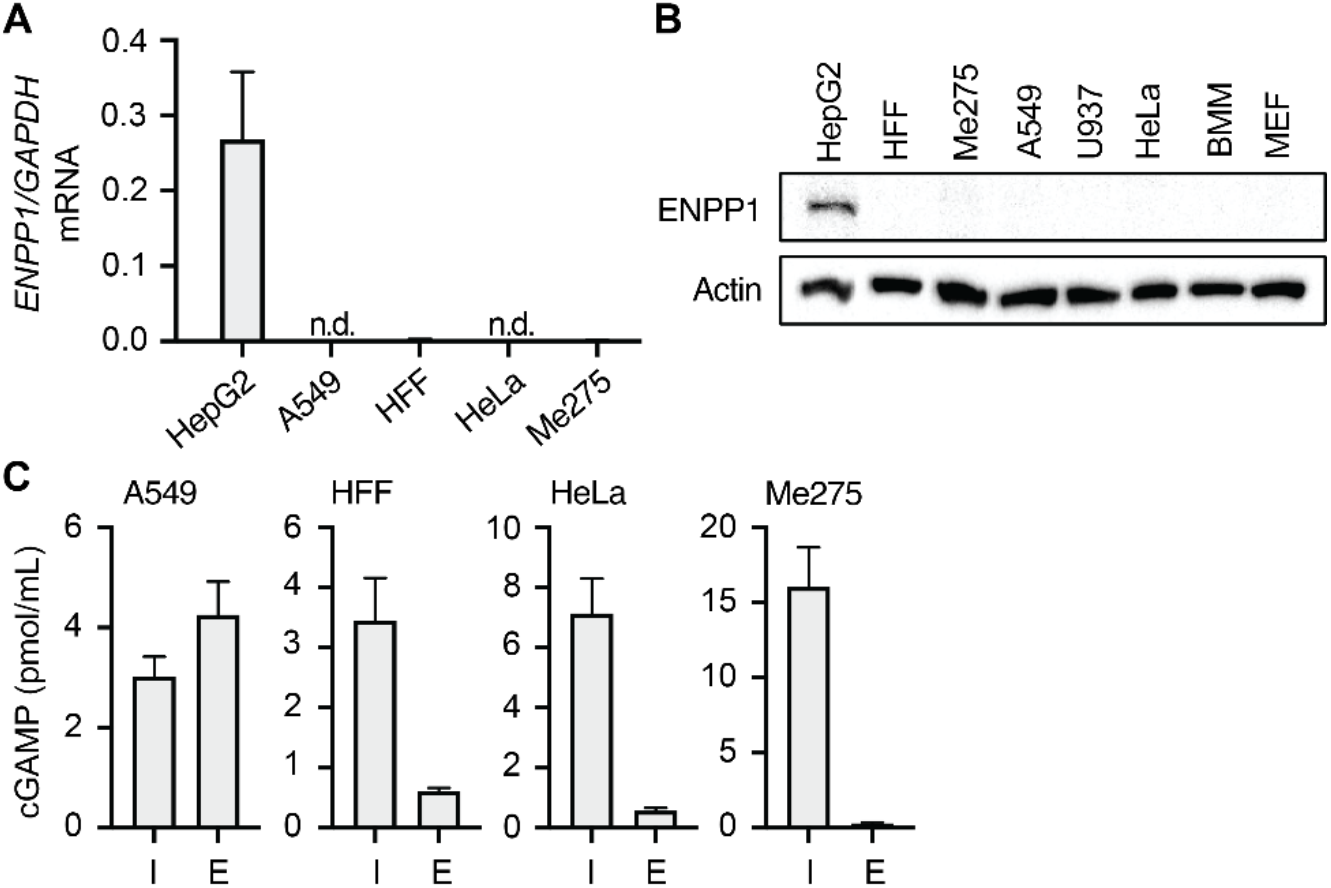
(**A**) Quantification of *ENPP1* mRNA transcript in indicated cell types by RT-qPCR. HepG2 cells are used as control for *ENPP1* expression. (**B**) Quantification of ENPP1 protein by Western blot. HepG2 cells are used as control for ENPP1 expression. Error bars represent mean ± SD. All data are representative of three independent experiments. (**C**) Quantification of intracellular and extracellular cGAMP from indicated cell types following CT DNA transfection for 8 hours.

**Fig. S2.**
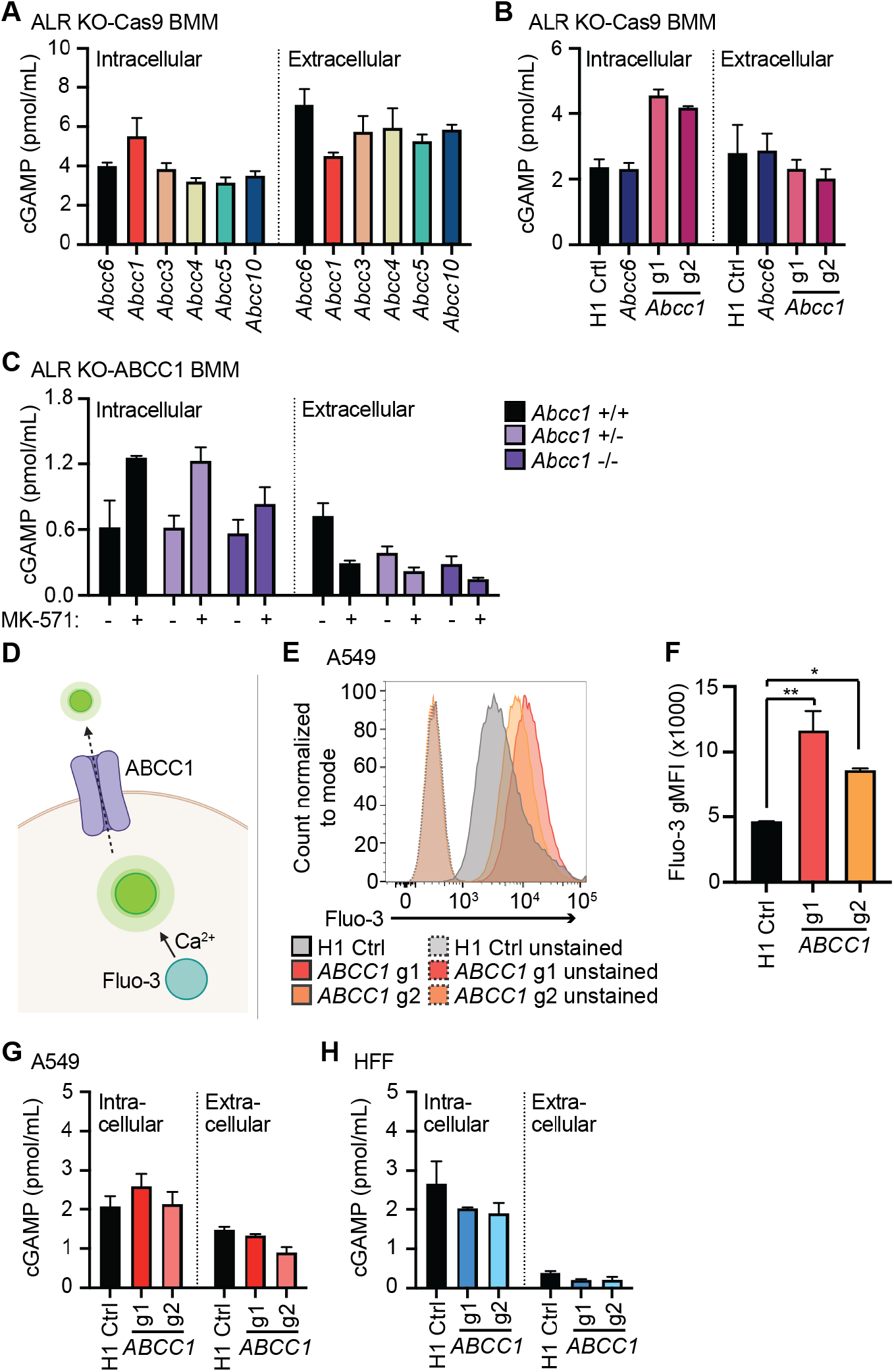
(**A, B**) Quantification of intracellular and extracellular cGAMP from indicated targeted lines of BMMs following CT DNA transfection for 8 hours. (**C**) Quantification of intracellular and extracellular cGAMP from indicated genotypes following CT DNA transfection for 8 hours. Cells were pretreated with 25 µM MK-571 or mock. (**D**) Schematic of Fluo-3AM staining and export. (**E**) *ABCC1*- or H1 control-targeted A549 cells were stained with Fluo-3AM and incubated for 1 hour at 37 C, followed by flow cytometric quantification of Fluo-3AM staining intensity. (**F**) gMFI quantification of flow cytometry data from (E). (**G, H**) Quantification of intracellular and extracellular cGAMP from indicated cell types following CT DNA transfection for 8 hours. Error bars represent mean ± SD. Statistical analysis was performed using a one-way ANOVA comparing all groups to H1 control and corrected for multiple comparisons using the Holm-Sidak method (E). *p<0.05, **p<0.01. All data are representative of two (C-E) or three (A, B, F, G) independent experiments.

**Fig. S3.**
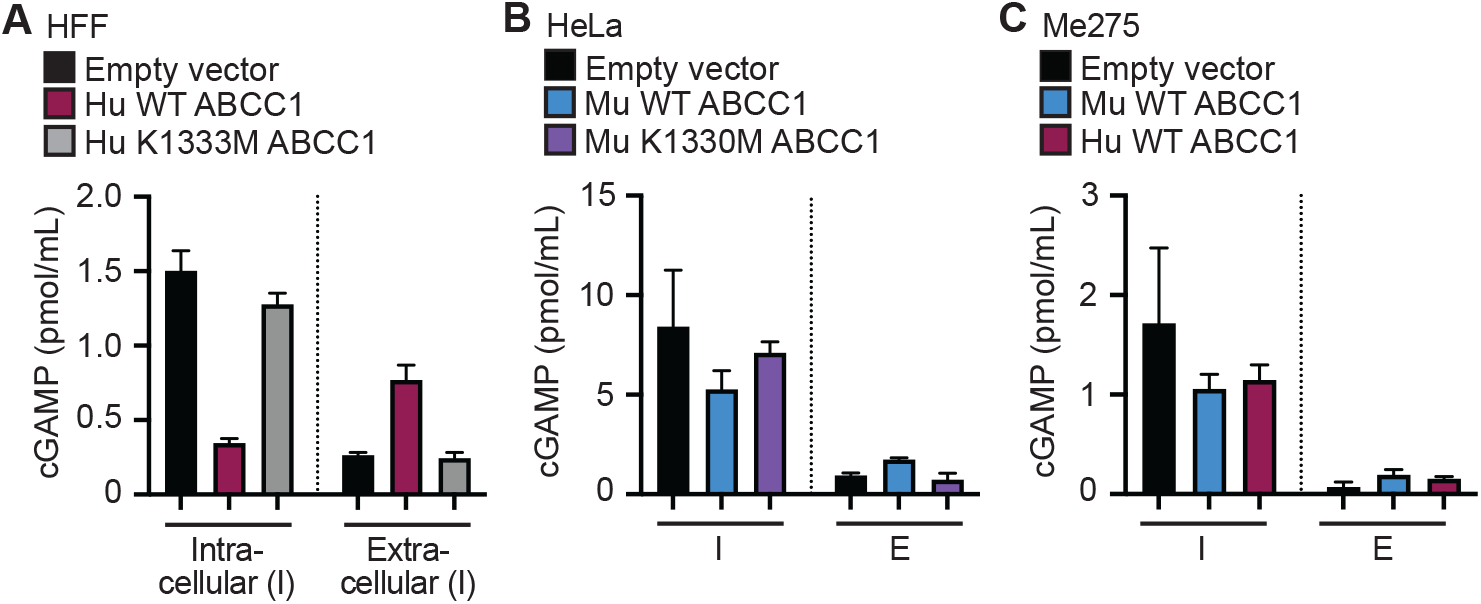
(**A**-**C**) Quantification of intracellular and extracellular cGAMP from indicated cell types following CT DNA transfection for 8 hours.

**Fig. S4.**
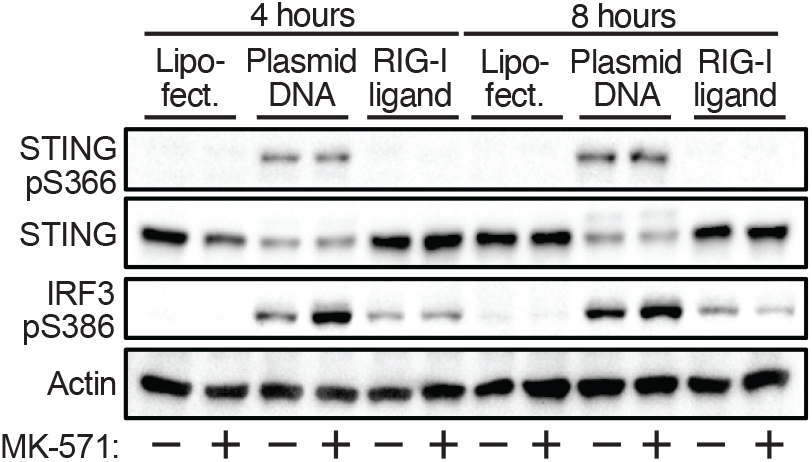
Western blot analysis of phosphorylated STING, STING, and phosphorylated IRF3 in HFFs that were treated with 25 µM MK-571 or mock followed by transfection with plasmid DNA or RIG-I ligand for 4 or 8 hrs.

## References

1. D. Goubau, S. Deddouche, C. Reise Sousa, Cytosolic sensing of viruses. Immunity 38, 855–869 (2013).

2. J. T. Crowl, E. E. Gray, K. Pestal, H. E. Volkman, D. B. Stetson, Intracellular Nucleic Acid Detection in Autoimmunity. Annu Rev Immunol 35, 313–336 (2017).

3. T. Li, Z. J. Chen, The cGAS-cGAMP-STING pathway connects DNA damage to inflammation, senescence, and cancer. J Exp Med 215, 1287–1299 (2018).

4. L. Sun, J. Wu, F. Du, X. Chen, Z. J. Chen, Cyclic GMP-AMP synthase is a cytosolic DNA sensor that activates the type I interferon pathway. Science 339, 786–791 (2013).

5. J. Wu et al., Cyclic GMP-AMP is an endogenous second messenger in innate immune signaling by cytosolic DNA. Science 339, 826–830 (2013).

6. H. Ishikawa, Z. Ma, G. N. Barber, STING regulates intracellular DNA-mediated, type I interferon-dependent innate immunity. Nature 461, 788–792 (2009).

7. S. Liu et al., Phosphorylation of innate immune adaptor proteins MAVS, STING, and TRIF induces IRF3 activation. Science 347, aaa2630 (2015).

8. J. J. Woodward, A. T. Iavarone, D. A. Portnoy, c-di-AMP secreted by intracellular Listeria monocytogenes activates a host type I interferon response. Science 328, 1703–1705 (2010).

9. P. J. Kranzusch et al., Ancient Origin of cGAS-STING Reveals Mechanism of Universal 2’,3’ cGAMP Signaling. Mol Cell 59, 891–903 (2015).

10. J. W. Nelson, R. R. Breaker, The lost language of the RNA World. Sci Signal 10, (2017).

11. L. Li et al., Hydrolysis of 2’3’-cGAMP by ENPP1 and design of nonhydrolyzable analogs. Nat Chem Biol 10, 1043–1048 (2014).

12. K. Kato et al., Structural insights into cGAMP degradation by Ecto-nucleotide pyrophosphatase phosphodiesterase 1. Nat Commun 9, 4424 (2018).

13. A. Ablasser et al., Cell intrinsic immunity spreads to bystander cells via the intercellular transfer of cGAMP. Nature 503, 530–534 (2013).

14. A. Bridgeman et al., Viruses transfer the antiviral second messenger cGAMP between cells. Science 349, 1228–1232 (2015).

15. M. Gentili et al., Transmission of innate immune signaling by packaging of cGAMP in viral particles. Science 349, 1232–1236 (2015).

16. A. Marcus et al., Tumor-Derived cGAMP Triggers a STING-Mediated Interferon Response in Non-tumor Cells to Activate the NK Cell Response. Immunity 49, 754-763.e754 (2018).

17. J. Carozza et al., Extracellular cGAMP is a cancer-cell-produced immunotransmitter involved in radiation-induced anticancer immunity. Nature Cancer 1, 184–196 (2020).

18. R. D. Luteijn et al., SLC19A1 transports immunoreactive cyclic dinucleotides. Nature 573, 434–438 (2019).

19. C. Ritchie, A. F. Cordova, G. T. Hess, M. C. Bassik, L. Li, SLC19A1 Is an Importer of the Immunotransmitter cGAMP. Mol Cell 75, 372-381.e375 (2019).

20. L. J. Lahey et al., LRRC8A:C/E Heteromeric Channels Are Ubiquitous Transporters of cGAMP. Mol Cell 80, 578-591.e575 (2020).

21. C. Zhou et al., Transfer of cGAMP into Bystander Cells via LRRC8 Volume-Regulated Anion Channels Augments STING-Mediated Interferon Responses and Anti-viral Immunity. Immunity 52, 767-781.e766 (2020).

22. A. F. Cordova, C. Ritchie, V. Böhnert, L. Li, Human SLC46A2 Is the Dominant cGAMP Importer in Extracellular cGAMP-Sensing Macrophages and Monocytes. ACS Cent Sci 7, 1073–1088 (2021).

23. Y. Zhou et al., Blockade of the Phagocytic Receptor MerTK on Tumor-Associated Macrophages Enhances P2X7R-Dependent STING Activation by Tumor-Derived cGAMP. Immunity 52, 357-373.e359 (2020).

24. V. Hornung et al., AIM2 recognizes cytosolic dsDNA and forms a caspase-1-activating inflammasome with ASC. Nature 458, 514–518 (2009).

25. S. L. Fink, B. T. Cookson, Apoptosis, pyroptosis, and necrosis: mechanistic description of dead and dying eukaryotic cells. Infect Immun 73, 1907–1916 (2005).

26. J. Maelfait, L. Liverpool, J. Rehwinkel, Nucleic Acid Sensors and Programmed Cell Death. J Mol Biol 432, 552–568 (2020).

27. E. E. Gray et al., The AIM2-like Receptors Are Dispensable for the Interferon Response to Intracellular DNA. Immunity 45, 255–266 (2016).

28. K. Burleigh et al., Human DNA-PK activates a STING-independent DNA sensing pathway. Sci Immunol 5, (2020).

29. H. E. Volkman, S. Cambier, E. E. Gray, D. B. Stetson, Tight nuclear tethering of cGAS is essential for preventing autoreactivity. Elife 8, (2019).

30. D. J. Giard et al., In vitro cultivation of human tumors: establishment of cell lines derived from a series of solid tumors. J Natl Cancer Inst 51, 1417–1423 (1973).

31. G. O. Gey, W. D. Coffman, M. T. Kubicek, Tissue culture studies of the proliferative capacity of cervical carcinoma and normal epithelium. Cancer Research 12, 264–265 (1952).

32. D. Valmori et al., Enhanced generation of specific tumor-reactive CTL in vitro by selected Melan-A/MART-1 immunodominant peptide analogues. J Immunol 160, 1750–1758 (1998).

33. M. Dean, A. Rzhetsky, R. Allikmets, The human ATP-binding cassette (ABC) transporter superfamily. Genome Res 11, 1156–1166 (2001).

34. V. Vasiliou, K. Vasiliou, D. W. Nebert, Human ATP-binding cassette (ABC) transporter family. Hum Genomics 3, 281–290 (2009).

35. R. W. Robey et al., Revisiting the role of ABC transporters in multidrug-resistant cancer. Nat Rev Cancer 18, 452–464 (2018).

36. Y. L. Sun, A. Patel, P. Kumar, Z. S. Chen, Role of ABC transporters in cancer chemotherapy. Chin J Cancer 31, 51–57 (2012).

37. A. R. Safa, Photoaffinity labeling of the multidrug-resistance-related P-glycoprotein with photoactive analogs of verapamil. Proc Natl Acad Sci U S A 85, 7187–7191 (1988).

38. J. D. Allen et al., Potent and specific inhibition of the breast cancer resistance protein multidrug transporter in vitro and in mouse intestine by a novel analogue of fumitremorgin C. Mol Cancer Ther 1, 417–425 (2002).

39. M. Dean, T. Annilo, Evolution of the ATP-binding cassette (ABC) transporter superfamily in vertebrates. Annu Rev Genomics Hum Genet 6, 123–142 (2005).

40. L. D. Weidner et al., The Inhibitor Ko143 Is Not Specific for ABCG2. J Pharmacol Exp Ther 354, 384–393 (2015).

41. P. Ralph, M. A. Moore, K. Nilsson, Lysozyme synthesis by established human and murine histiocytic lymphoma cell lines. J Exp Med 143, 1528–1533 (1976).

42. I. Leier et al., The MRP gene encodes an ATP-dependent export pump for leukotriene C4 and structurally related conjugates. J Biol Chem 269, 27807–27810 (1994).

43. S. P. Cole, Multidrug resistance protein 1 (MRP1, ABCC1), a “multitasking” ATP-binding cassette (ABC) transporter. J Biol Chem 289, 30880–30888 (2014).

44. P. Mitra et al., Role of ABCC1 in export of sphingosine-1-phosphate from mast cells. Proc Natl Acad Sci U S A 103, 16394–16399 (2006).

45. G. Reid et al., Characterization of the transport of nucleoside analog drugs by the human multidrug resistance proteins MRP4 and MRP5. Mol Pharmacol 63, 1094–1103 (2003).

46. R. D. Barrington, P. W. Needs, G. Williamson, P. A. Kroon, MK571 inhibits phase-2 conjugation of flavonols by Caco-2/TC7 cells, but does not specifically inhibit their apical efflux. Biochem Pharmacol 95, 193–200 (2015).

47. R. J. Platt et al., CRISPR-Cas9 knockin mice for genome editing and cancer modeling. Cell 159, 440–455 (2014).

48. R. L. Brunette et al., Extensive evolutionary and functional diversity among mammalian AIM2-like receptors. J Exp Med 209, 1969–1983 (2012).

49. E. K. Brinkman, T. Chen, M. Amendola, B. van Steensel, Easy quantitative assessment of genome editing by sequence trace decomposition. Nucleic Acids Res 42, e168 (2014).

50. D. Keppler, Y. Cui, J. König, I. Leier, A. Nies, Export pumps for anionic conjugates encoded by MRP genes. Adv Enzyme Regul 39, 237–246 (1999).

51. S. Prechtl et al., The multidrug resistance protein 1: a functionally important activation marker for murine Th1 cells. J Immunol 164, 754–761 (2000).

52. M. Gao et al., Comparison of the functional characteristics of the nucleotide binding domains of multidrug resistance protein 1. J Biol Chem 275, 13098–13108 (2000).

53. Y. J. Crow et al., Mutations in the gene encoding the 3’-5’ DNA exonuclease TREX1 cause Aicardi-Goutières syndrome at the AGS1 locus. Nat Genet 38, 917–920 (2006).

54. D. B. Stetson, J. S. Ko, T. Heidmann, R. Medzhitov, Trex1 prevents cell-intrinsic initiation of autoimmunity. Cell 134, 587–598 (2008).

55. E. E. Gray, P. M. Treuting, J. J. Woodward, D. B. Stetson, Cutting Edge: cGAS Is Required for Lethal Autoimmune Disease in the Trex1-Deficient Mouse Model of Aicardi-Goutières Syndrome. J Immunol 195, 1939–1943 (2015).

56. D. Gao et al., Activation of cyclic GMP-AMP synthase by self-DNA causes autoimmune diseases. Proc Natl Acad Sci U S A 112, E5699–5705 (2015).

57. A. Gall et al., Autoimmunity initiates in nonhematopoietic cells and progresses via lymphocytes in an interferon-dependent autoimmune disease. Immunity 36, 120–131 (2012).

58. A. Lorico et al., Disruption of the murine MRP (multidrug resistance protein) gene leads to increased sensitivity to etoposide (VP-16) and increased levels of glutathione. Cancer Res 57, 5238–5242 (1997).

59. P. R. Wielinga et al., Characterization of the MRP4-and MRP5-mediated transport of cyclic nucleotides from intact cells. J Biol Chem 278, 17664–17671 (2003).

60. G. Jedlitschky, B. Burchell, D. Keppler, The multidrug resistance protein 5 functions as an ATP-dependent export pump for cyclic nucleotides. J Biol Chem 275, 30069–30074 (2000).

61. Y. Guo et al., MRP8, ATP-binding cassette C11 (ABCC11), is a cyclic nucleotide efflux pump and a resistance factor for fluoropyrimidines 2’,3’-dideoxycytidine and 9’-(2’-phosphonylmethoxyethyl)adenine. J Biol Chem 278, 29509–29514 (2003).

62. A. T. Whiteley et al., Bacterial cGAS-like enzymes synthesize diverse nucleotide signals. Nature 567, 194–199 (2019).

63. B. Lowey et al., CBASS Immunity Uses CARF-Related Effectors to Sense 3’-5’-and 2’-5’-Linked Cyclic Oligonucleotide Signals and Protect Bacteria from Phage Infection. Cell 182, 38-49.e17 (2020).

64. X. Gui et al., Autophagy induction via STING trafficking is a primordial function of the cGAS pathway. Nature 567, 262–266 (2019).

65. B. R. Morehouse et al., STING cyclic dinucleotide sensing originated in bacteria. Nature 586, 429–433 (2020).

66. A. M. Burroughs, D. Zhang, D. E. Schäffer, L. M. Iyer, L. Aravind, Comparative genomic analyses reveal a vast, novel network of nucleotide-centric systems in biological conflicts, immunity and signaling. Nucleic Acids Res 43, 10633–10654 (2015).

67. C. F. Higgins, ABC transporters: from microorganisms to man. Annu Rev Cell Biol 8, 67–113 (1992).

68. S. Orozco et al., RIPK1 both positively and negatively regulates RIPK3 oligomerization and necroptosis. Cell Death Differ 21, 1511–1521 (2014).

69. T. Saito, D. M. Owen, F. Jiang, J. Marcotrigiano, M. Gale, Innate immunity induced by composition-dependent RIG-I recognition of hepatitis C virus RNA. Nature 454, 523–527 (2008).

70. N. E. Sanjana, O. Shalem, F. Zhang, Improved vectors and genome-wide libraries for CRISPR screening. Nat Methods 11, 783–784 (2014).

